# *In vitro* efficacy of gentamicin against multidrug-resistant *Neisseria gonorrhoeae*: synergy of three gentamicin antimicrobial combinations

**DOI:** 10.1101/2021.04.26.441563

**Authors:** Xuechun Li, Wenjing Le, Xiangdi Lou, Biwei Wang, Caroline A. Genco, Peter A. Rice, Xiaohong Su

## Abstract

**Objectives:** To determine *in vitro* activities of gentamicin alone and in combination with ceftriaxone, ertapenem and azithromycin against multidrug-resistant (MDR) *N. gonorrhoeae* isolates.

**Methods:** 407 isolates from Nanjing, China, obtained in 2016 and 2017, had minimum inhibitory concentrations (MICs) determined for gentamicin using the agar dilution method. Antimicrobial combinations were also tested in 97 MDR strains using the antimicrobial gradient epsilometer test (Etest); results ranging from synergy to antagonism were interpreted using the fractional inhibitory concentration (FICI).

**Results:** All 407 gonococcal isolates were susceptible to gentamicin. MICs ranged from 2 mg/L to 16 mg/L. Synergy was demonstrated in 16.5%(16/97), 27.8%(27/97) and 8.2%(8/97) MDR strains when gentamicin was combined with ceftriaxone [geometric mean (GM) FICI; 0.747], ertapenem (GM FICI; 0.662) and azithromycin (GM FICI; 1.021), respectively. No antimicrobial antagonism was observed with any combination. The three antimicrobial combinations were indifferent overall. The overall GM MICs of gentamicin were reduced by 2.63-, 3.80- and 1.98-fold when tested in combination with ceftriaxone, ertapenem and azithromycin, respectively. The GM MICs of the three antimicrobials by themselves were reduced by 3-, 2.57- and 1.98-fold respectively, when each was tested in combination with gentamicin. No antimicrobial antagonism was observed with any combination.

**Conclusions:** Gentamicin alone was effective *in vitro* against MDR *N. gonorrhoeae* and in combination with ceftriaxone, ertapenem or azithromycin. Combination testing of resistant strains, overall, showed lower effective MICs against gentamicin itself and each of the three antimicrobials when used in combination with gentamicin.

## INTRODUCTION

Gonorrhea, caused by *Neisseria gonorrhoeae*, is a common bacterial sexually transmitted infection (STI), which is responsible for an increasing burden of disease globally. In 2016, the World Health Organization (WHO) estimated 87 million new cases of gonorrhea worldwide in adults, aged 15-49(1). Common features of gonococcal infection include urethritis and cervicitis; more serious manifestations include pelvic inflammatory disease, chronic pelvic pain, infertility, chorioamnionitis in women and epididymitis in men. Disseminated gonococcal infections (DGI) occurs occasionally in adults and neonates born to infected mothers; in the pre-antibiotic era endocarditis and meningitis sometimes resulted from gonococcal bacteremia but is rarely seen today. Concomitant gonococcal and HIV infections also increase the risk of HIV transmission (2).

The emergence of antibiotic resistance among *N. gonorrhoeae* is a global public health threat. *N. gonorrhoeae* has developed resistance to all antimicrobials that have been used for treatment, including penicillin, tetracycline, ciprofloxacin and azithromycin, leading to the emergence of multidrug-resistant (MDR) isolates, which are difficult to treat (3). Currently, dual antimicrobial therapy with an extended spectrum cephalosporin (ESC), either ceftriaxone 250 mg intramuscularly or cefixime 400 mg orally, plus azithromycin 1g orally, is recommended as first-line treatment of uncomplicated gonorrhea by the World Health Organization (WHO) (4). Increased prevalence of azithromycin resistance globally prompted a revision of prior recommendations in the United Kingdom (U.K.) (5) in 2018 and the United States (U.S.) (6) in 2020 from dual therapy with ceftriaxone and azithromycin to recommendations of higher doses of ceftriaxone: 1 gram (U.K.) and 500 mg (U.S). Unfortunately, resistance to ESCs (7-9), the last remaining option for empirical first-line monotherapy, threatens future use of this class of antimicrobials. Treatment failures with both mono- and dual-therapy (including azithromycin) have been reported in recent years (10, 11).

Treatment options for gonorrhea including infections caused by MDR organisms are diminishing; there is an urgent need to explore new or repurposed antimicrobial agents and/or therapeutic strategies. Gentamicin, an aminoglycoside antibiotic that inhibits protein synthesis by binding to the 30S ribosomal subunit, has been used as first-line therapy for treatment of gonorrhea in several countries, including Malawi, where it has been used officially for nearly 30 years (12). Numerous *in vitro* susceptibility studies have shown that gentamicin is active against *N. gonorrhoeae* (13-16), suggesting that gentamicin is appropriate for treatment of uncomplicated gonococcal infection. A single 240 mg dose of intramuscular gentamicin was 95% efficacious in a randomized controlled clinical trial to treat urethritis (17). In multicenter clinical trials, dual therapy with gentamicin plus azithromycin was 94%-100% effective compared to 98% for ceftriaxone plus azithromycin (18, 19). Little is known about *in vitro* susceptibility of gentamicin in Chinese isolates where this antimicrobial has not been used to treat gonorrhea. Use of antimicrobial combinations is a therapeutic strategy intended to increase efficacy and slow the development of resistance (20). Antimicrobial combinations may prove useful to successfully manage MDR *N. gonorrhoeae* infections and they are included in current WHO and CDC guidelines (4, 6).

Initially, we evaluated gentamicin susceptibility of gonococcal strains isolated between 2016 and 2017 in Nanjing, China. Second, we carried out studies with antimicrobial combinations that included gentamicin to evaluate possible *in vitro* enhancement of gentamicin activity against MDR strains when tested in combination with either ceftriaxone (an ESC), ertapenem (a carbapenem) or azithromycin (a macrolide). Third, we determined the MICs of each of the three antimicrobials by themselves when tested in combination with gentamicin.

## RESULTS

All 407 *N. gonorrhoeae* strains were sensitive to gentamicin by agar dilution; minimum inhibitory concentrations (MICs) ranged from 2mg/L to 16mg/L; the MIC_50_ was 8 mg/L and the MIC_90_ was 16mg/L. Among the 407 isolates, 34 (8.4%) were susceptible (MIC ≤ 4mg/L), 373 (91.6%) were intermediately susceptible (MIC range, 8-16 mg/L) and none was resistant (MIC ≥ 32 mg/L) (Fig.1). Among the 34 fully susceptible isolates, 6 (17.6%) had decreased susceptibility to ESCs i.e. ceftriaxone or cefixime or both and 1 (2.9%) was fully resistant to ceftriaxone (MIC; 1mg/L) and cefixime (MIC; 2mg/L). There was no change in intermediate susceptibility to gentamicin of isolates between 2016 and 2017 (89.3% vs. 93.4%; χ^2^=2.225, P=0.136) (Fig. 2).

**Figure 1.**
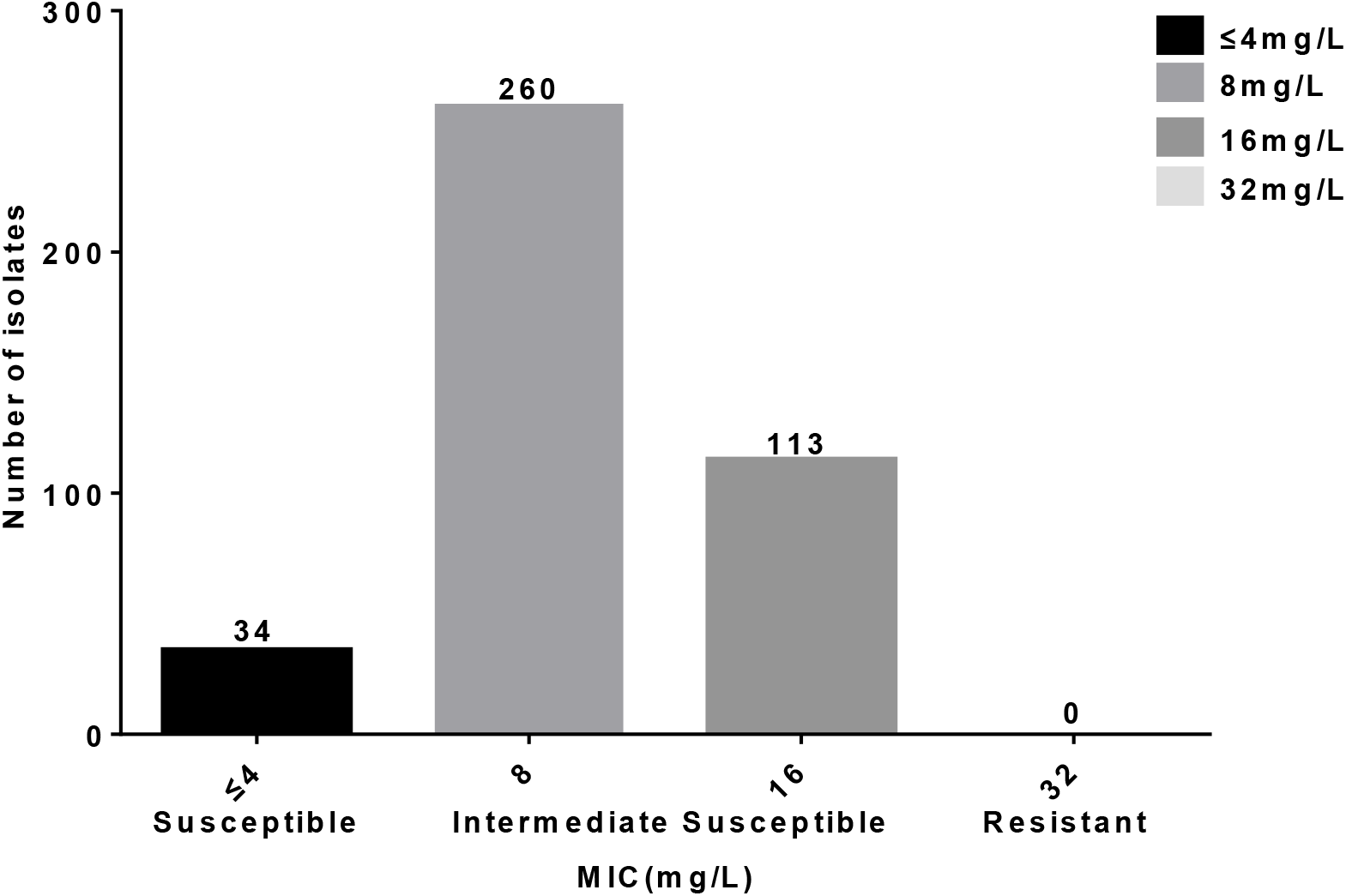
Susceptibility of gentamicin against *N. gonorrhoeae* strains isolated in Nanjing, China, 2016–2017 (n =407).

**Figure 2.**
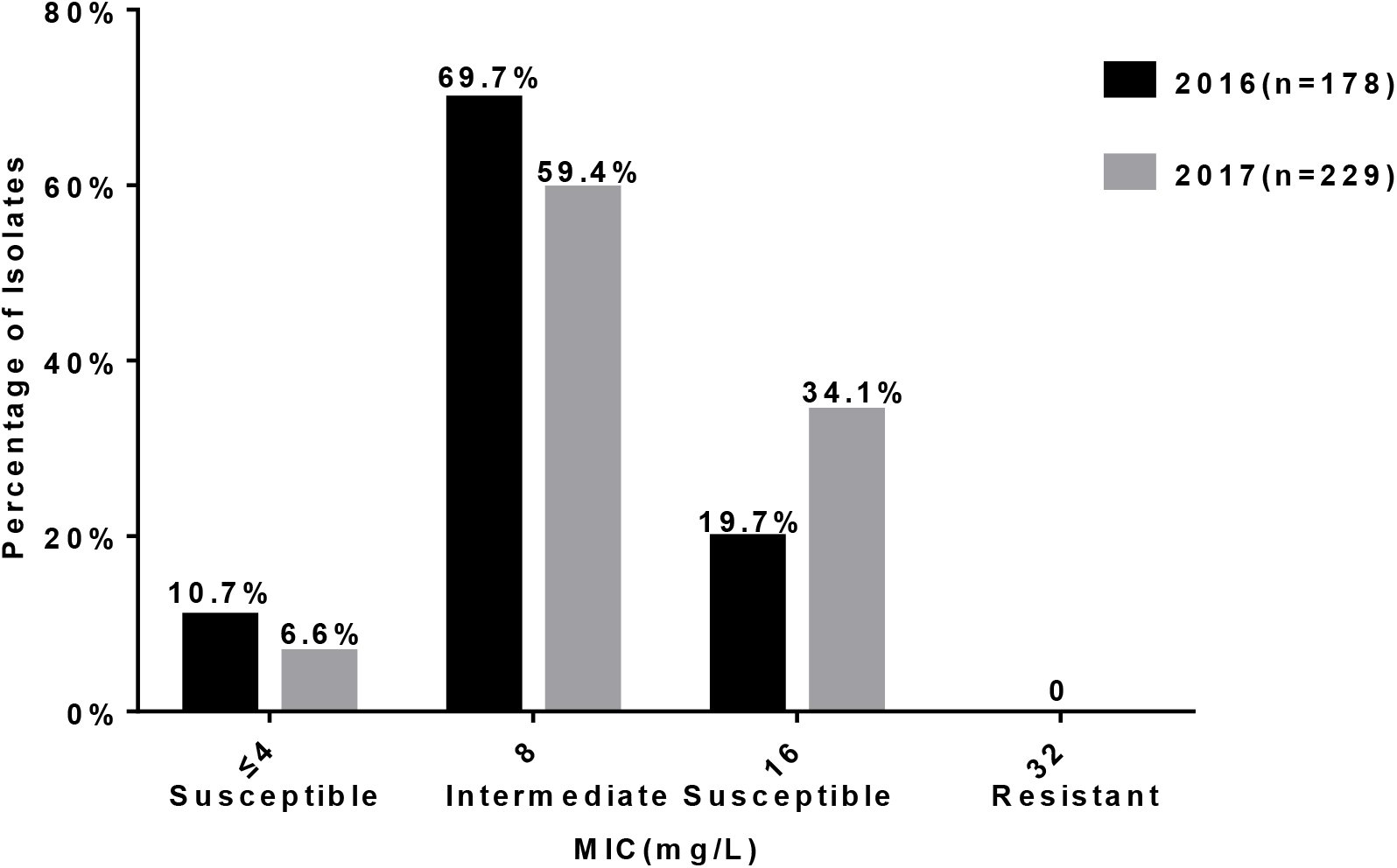
Distributions of MICs of gentamicin against *N. gonorrhoeae* isolates in 2016 (n=178) and 2017 (n=229).

The distribution of antimicrobial susceptibility patterns for the 97 MDR strains selected for antimicrobial combination testing showed 6 unique patterns of susceptibility (Table S1). 93 (95.9%) strains had decreased susceptibility to ceftriaxone or cefixime or both plus resistance to ciprofloxacin and penicillin, and 4 (4.1%) strains had decreased susceptibility to ceftriaxone or cefixime or both plus resistance to ciprofloxacin, penicillin and azithromycin. Resistance to gentamicin in MDR isolates was not detected by either the agar dilution or the Etest method. Agreement of MICs between agar dilution and Etest among MDR isolates is summarized in Table S2. Overall agreement of MICs (≤ 2-fold different) between the two methods was 93.8%. Etest always resulted in one to two dilutions lower MIC values than agar dilution.

Gentamicin agar dilution versus Etest MIC results (mg/L) were as follows: MIC_50_ = 8 versus 6; MIC_90_ =16 versus 8; geometric mean [GM] MIC =11.3 versus 5.84. Additionally, there was a discrepancy in the full susceptibility category (29.9% fully susceptible by Etest versus 9.3% by agar dilution; χ^2^=13.090, P<0.001).

The three antimicrobial combinations used to test each of the 97 strains were examined for effects that were classified as: synergistic, indifferent or antagonistic--summarized in Table 1. For example, the gentamicin GM MIC, when tested alone, was 5.840 mg/L; when combined with ceftriaxone, the GM MIC was reduced to 2.217 mg/L (2.63-fold reduction, P <0.001) (Table 2). Together with ceftriaxone, gentamicin exhibited synergy against 16.5% (16/97) MDR strains; overall, the combination was indifferent; FICI, 0.747. When tested alone, MICs of ceftriaxone ranged from 0.016-0.75 mg/L; in combination with gentamicin, the GM MIC against ceftriaxone decreased from 0.078 to 0.026 mg/L (3-fold reduction, P <0.001).

**Table 1.** Synergy test results for combinations of gentamicin plus azithromycin, ceftriaxone, ertapenem against 97 MDR *N. gonorrhoeae* isolates

| Effect n (%) | Antimicrobial combinations |  |  |
| --- | --- | --- | --- |
|  | GEN +CRO | GEN +ETP | GEN +AZM |
| Synergistic n (%) | 16(16.5%) | 27(27.8%) | 8(8.2%) |
| Indifferent n (%) | 81(83.5%) | 70(72.2%) | 89(91.8%) |
| Antagonistic n (%) | 0 | 0 | 0 |
| FICI (geometric mean) | 0.747 | 0.662 | 1.021 |
| Classification Overall | Indifferent | Indifferent | Indifferent |

Abbreviations: n, number; GEN, gentamicin; CRO, ceftriaxone; ETP, ertapenem;
AZM, azithromycin.

**Table 2.** *In vitro* antibacterial activity of tested antibiotics combinations against 97 MDR *N. gonorrhoeae* isolates

| MIC | Antimicrobial combinations |  |  |
| --- | --- | --- | --- |
|  | GEN +CRO | GEN +ETP | GEN +AZM |
| MIC A<sub>alone</sub> | 5.840(2-12) | 5.840(2-12) | 5.840(2-12) |
| MIC A<sub>in combination</sub> | 2.217(0.38-6) | 1.536(0.25-6) | 2.949(1-8) |
| MIC B<sub>alone</sub> | 0.078(0.016-0.75) | 0.018(0.006-0.064) | 0.347(0.047-8) |
| MIC B<sub>in combination</sub> | 0.026(0.004-0.25) | 0.007(0.002-0.047) | 0.175(0.023-2) |

Abbreviations: GEN, gentamicin; CRO, ceftriaxone; ETP, ertapenem; AZM,
azithromycin; MIC, minimum inhibitory concentration.

The gentamicin GM MIC when combined with ertapenem decreased to 1.536 mg/L (3.80-fold reduction, P <0.001) (Table 2). Gentamicin together with ertapenem was the most synergistic combination, displaying synergy against 27.8% (27/97) of MDR gonococcal isolates; overall, this combination was indifferent; FICI, 0.662. Ertapenem MICs of 97 MDR isolates, when tested alone, ranged from 0.006 to 0.064 mg/L (GM MIC, 0.018 mg/L); when combined with gentamicin, the GM MIC decreased to 0.007 mg/L (2.57-fold reduction, P <0.001).

The gentamicin GM MIC when combined with azithromycin decreased to 2.949 mg/L (1.98-fold reduction, P <0.001) (Table 2). Together with azithromycin, gentamicin exhibited synergy against 8.2% (8/97) MDR strains; overall, the combination was indifferent; FICI, 1.021. When tested alone, MICs of azithromycin ranged from 0.047-8 mg/L; in combination with gentamicin, the GM MIC against azithromycin decreased from 0.347 to 0.175 mg/L (1.98-fold reduction, P <0.001).

## DISCUSSION

Our study provides data on gentamicin susceptibility against *N. gonorrhoeae* isolated in Nanjing (Jiangsu Province), China. On agar dilution testing, most strains displayed intermediate susceptibility to gentamicin (MIC 8-16 mg/L), similar to a study that examined gonococcal isolates from seven hospitals in a neighboring eastern Chinese province; in that study 97.8% (493/504) of strains possessed gentamicin MICs of 8-16 mg/L (21). European and U.S. studies have reported 82.7% (15) and 73% (18), intermediate susceptibility, respectively, of *N. gonorrhoeae* isolates to gentamicin.

A recent report of gentamicin susceptibility of *N. gonorrhoeae* in which 86.0% of 470 isolates were fully susceptible (MICs ≤ 4 mg/L), examined isolates from seven geographically distributed Chinese provinces as part of the China Gonococcal Resistance Surveillance Programme (China-GRSP) (22) similar to an Indian study where 90.7% of isolates were reported as fully susceptible (15).

Our study compared gentamicin MICs using agar dilution and Etest methods for 97 multidrug-resistant (MDR) *N. gonorrhoeae* isolates. Similar to previous studies (13, 23), we found that over 90% of gentamicin MICs determined by agar dilution and Etest were ≤ 2-fold different; typically, Etest resulted in lower MICs and identified a larger proportion of fully susceptible isolates. In particular, all MDR isolates were fully or intermediately susceptible to gentamicin.

Synergistic or additive effects of combining antimicrobials for treatment may slow the development of antimicrobial resistance of *N. gonorrhoeae* (24). We assessed the *in vitro* activity of gentamicin in combination with 3 antibiotics. Determining synergy has several challenges. Several test methods are available to evaluate synergistic effects of antimicrobial combinations, however, they are not well standardized. We chose Etest because it is practical and also correlates well with agar dilution, time–kill curves and checkerboard testing in demonstrating synergy for two-drug combination (25-27). Nonetheless, FICI interpretation depends on the criteria used. FICI values between 0.5 and 1 have been interpreted as additive in Indian (28) and Japanese (29) studies, differing from our criteria, which classifies FICI values in this range as indifferent.

Ceftriaxone, in higher doses, is now recommended as single therapy by the U.S. and the U.K. for treatment of uncomplicated gonorrhea (5, 6). Our study showed that the combination of ceftriaxone and gentamicin exhibited an indifferent effect overall in >80% of MDR strains (< 20% synergy). In the Indian study by Singh *et al* (28), 14.7% synergy and 6.3% antagonism were reported for this combination against 95 *N. gonorrhoeae* strains including 79 MDR and one extensively drug-resistant (XDR) strain. In a Canadian study, a mean FICI_50_ value of 1.2 (0.8-2.0) was shown for nine reference strains of *N. gonorrhoeae* (WHO F, G, K, L, M, N, O, P and ATCC 49226) with this combination (30). An American study reported a mean FICI of 1.25 (0.73-2) using gonococcal isolates that displayed different cefixime MICs (26). No synergistic/antagonistic effect (resulting in 100% indifference) was observed in either study (26, 30).

We chose ertapenem as a candidate for *in vitro* synergy testing because its mechanism of action differs from that of gentamicin and it has been used to treat infection with combined high-level azithromycin and ceftriaxone resistant *N. gonorrhoeae* (11). Ertapenem has demonstrated an advantage over ceftriaxone for MDR or ceftriaxone resistant isolates and has also been suggested for possible use in a dual antimicrobial regimen (31). We showed that gentamicin plus ertapenem in combination resulted in synergistic and indifferent effects with no antagonism demonstrated in any MDR strain. In the study by Singh *et al* (28), this combination displayed either synergy (31.6%) or indifference (68.4%) in 100% of strains; no antagonism was seen. Gentamicin in combination with azithromycin is currently recommended as an option for retreatment by the WHO when dual therapy fails (4). Also, it is proposed as an alternative CDC recommendation when higher dose ceftriaxone therapy cannot be used (6). In our studies, this combination demonstrated synergy in fewer MDR isolates (<10%) than combinations with either ceftriaxone or ertapenem and exhibited the highest FICI value of the three combinations tested. Similar to our results, Sood *et al* (32) demonstrated synergistic effects in 22.9% of isolates displaying different ceftriaxone MICs and no antagonism for this combination. All isolates with differing cefixime/ ceftriaxone MICs showed indifference with a mean FICI of 1.7/0.83 in studies from the UK (33) and Japan (29). The separate study by Singh *et al* (28) differed from these results, with 6.3% of strains exhibiting antagonism when this combination was used.

A summary of results from these previous studies is shown in Table S3. No synergistic/antagonistic effect (resulting in 100% indifference) was observed in studies from Canada (30), Japan (29), the US (26) and the UK (33). None of these studies incorporated isolates with multi-drug resistance. However, a certain proportion of synergistic effects was observed in the two Indian studies (28, 32). Antagonism was also observed in combinations of gentamicin with ceftriaxone and azithromycin in the study by Singh *et al* (28). Sood *et al* (32) used the same Etest that we used. In contrast, Singh *et al* (28) incubated the Etest strip of the first antimicrobial (antimicrobial A) for 1 h, and then replaced it with the Etest strip of the second antimicrobial (antimicrobial B) at the same location and looked for synergism. A mild degree of antagonism may have been missed in our tests when the zone of inhibition ran under the strips where they crossed and therefore was unreadable and interpreted as indifference (25).

From a pharmacokinetic perspective, all antimicrobials in the 3 combinations in our study result in peak levels of drug during the first 3 hours after administration (34-37). However, azithromycin has a longer half-life than gentamicin (approximately 68 and 2 hours, respectively) (34, 37). Gentamicin in combination with azithromycin also produced the lowest synergistic effects among the 3 combinations in our study so it may not be optimal for clinical use where synergy would not be prolonged. In conclusion, resistance to gentamicin was not observed in gonococcal isolates examined in this study, including MDR isolates, indicating potential for its therapeutic use. Antimicrobial combinations of gentamicin plus ertapenem, ceftriaxone, and azithromycin showed no antagonistic effects; enhanced efficacy of individual antimicrobials in the presence of other antimicrobials was also demonstrated. Improved efficacy of antimicrobial combinations suggest that dual therapy might be a promising approach to manage MDR *N. gonorrhoeae*. Further studies to correlate *in vitro* results with clinical outcomes are warranted.

## MATERIALS AND METHODS

### Bacterial strains

407 gonococcal isolates were recovered from men with symptomatic urethritis (urethral discharge and/or dysuria) attending the STD clinic at the Institute of Dermatology, Chinese Academy of Medical Sciences in Nanjing, between January 2016 and December 2017. Urethral specimens were collected with cotton swabs and immediately streaked onto modified Thayer-Martin medium (Zhuhai DL Biotech Co. Ltd.) and cultured in candle jars at 36°C for 24–48 h. Gonococcal isolates were identified by colonial morphology, Gram’s stain, and oxidase testing and sub-cultured onto GC chocolate agar base (Difco, Detroit, MI) supplemented with 1% IsovitaleX™ (Oxoid, USA); pure cultures were swabbed, suspended in tryptone-based soy broth and frozen (™80°C) until used for antimicrobial testing.

### Antimicrobia l susceptibility testing

The MICs of the 407 isolates were determined using the agar dilution method for gentamicin, penicillin, tetracycline, ciprofloxacin, azithromycin, spectinomycin, cefixime and ceftriaxone, used singly, according to the Clinical and Laboratory Standards Institute (CLSI) guidelines (38). *N. gonorrhoeae* ATCC 49226, WHO reference strains F, G, L, O and P were used as quality control strains in susceptibility tests. Although formal susceptibility criteria for gentamicin have not been established by CLSI, criteria described by CDC (16) have used MICs of ≤4 mg/L as fully susceptible, 8 to 16 mg/L as intermediately susceptible and ≥32 mg/L as resistant. Resistance to azithromycin (MIC ≥1 mg/L) was determined using EUCAST criteria (39). Susceptibilities to other antibiotics were assessed based on CLSI standards (38). Decreased susceptibility to cephalosporins was determined according to WHO standards (40). Based on criteria proposed by Tapsall *et al* in 2009 (41), MDR isolates were defined as those resistant or with decreased susceptibility to one or more widely used antimicrobials (ceftriaxone and cefixime; category I) and resistant to two or more antimicrobials, which are used less frequently (penicillin; ciprofloxacin and azithromycin; category II) (Table S4).

### Synergy testing and interpretation

Of 407 clinical isolates of *N. gonorrhoeae*, 97 MDR isolates (determined below) were selected for antimicrobial combination testing (synergy/ antagonism) according to MDR criteria. WHO-P was used as a quality control strain. Dual antimicrobial testing was performed to evaluate the efficacy of gentamicin in combination with either ceftriaxone, ertapenem or azithromycin, using the Etest method, described previously (25). Briefly, MICs for individual antimicrobials (MIC A _alone_ and MIC B _alone_) against *Neisseria gonorrhoeae* isolates were determined using Etest strips (Liofilchem, Italy); *in vitro* activity of each combination was determined by placing E-test strips of the two antimicrobials on the agar plates at a 90° angle, with intersections at the points of their individual MICs (Fig. 3). Agar plates were inverted during incubation at 36°C in 5% CO_2_ for 16–18 hours, and the MIC of each antimicrobial in the combination (MIC A _in combination_ and MIC B _in combination_) was read.

**Figure 3.**
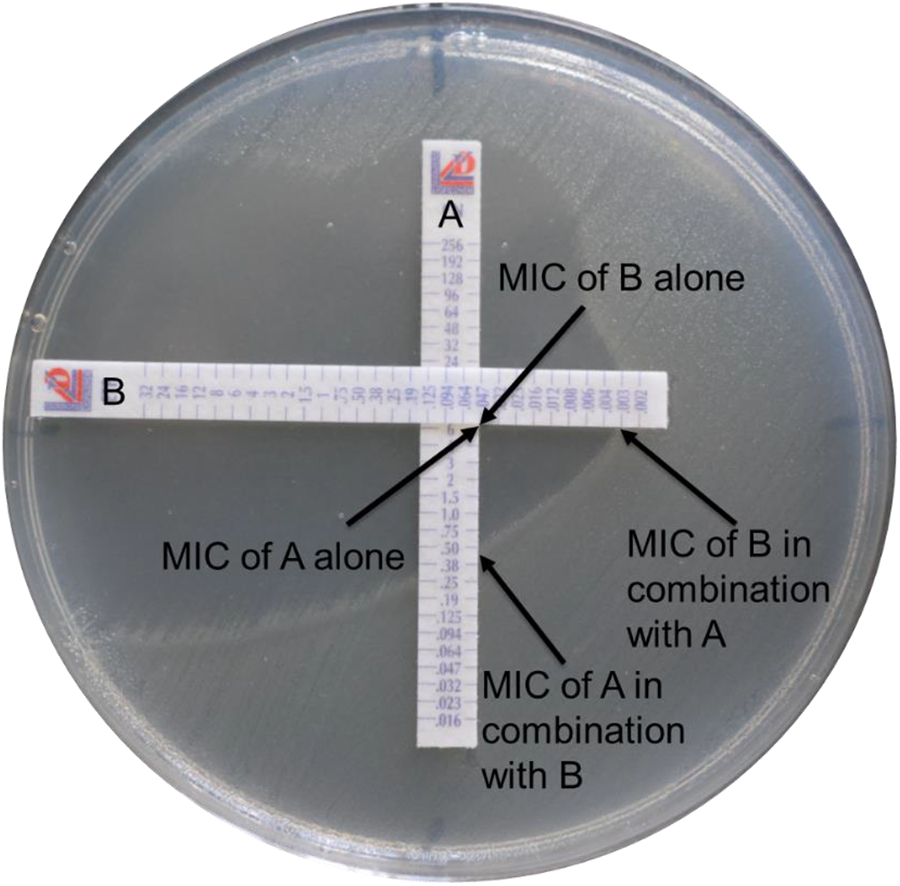
Photograph of strip placement for E-test synergy method.

To determine whether each antimicrobial combination resulted in a synergistic, indifferent or antagonistic effect, the fractional inhibitory concentration index (FICI) was calculated using the following formula: FICI = (MIC A _in combination_ / MIC A _alone_) +(MIC B _in combination_ / MIC B _alone_) (25). FICI values were interpreted using the following criteria: synergy: ≤0.5; indifference: FIC >0.5 to ≤4.0 and antagonism: FICI of > 4.0(25).

### Statistical Analysis

The chi-square test was used to compare gentamicin susceptibility trend data and the categorical assignments of gentamicin susceptibility by the agar dilution or Etest method. Mean values of MICs and FICIs were calculated as geometric means (GM). The statistical significance of the difference between the MIC of each of the three antimicrobials tested alone and in combination with gentamicin was determined using the nonparametric Mann-Whitney’s U-test. P values less than 0.05 were considered statistically significant.

## ACKNOWLEDGEMENTS

This work was supported by the grants from the Chinese Academy of Medical Sciences Initiative for Innovative Medicine (2016-I2M-3-021) and the U.S. National Institutes of Health (AI084048 and AI116969).

## CONFLICTS OF INTEREST

No conflicts for all authors.

## REFERENCES

1. World Health Organization. Report on global sexually transmitted infection surveillance -2018[M]. World Health Organization, 2018; Available from: https://www.who.int/publications/i/item/9789241565691.

2. Mlisana, K., N. Naicker, L. Werner, L. Roberts, F. van Loggerenberg, C. Baxter, J. A. Passmore, A. C. Grobler, A. W. Sturm, C. Williamson, K. Ronacher, G. Walzl, and K. S. Abdool. 2012. Symptomatic vaginal discharge is a poor predictor of sexually transmitted infections and genital tract inflammation in high-risk women in South Africa. J Infect Dis 206:6–14.

3. Unemo, M., C. S. Bradshaw, J. S. Hocking, H. de Vries, S. C. Francis, D. Mabey, J. M. Marrazzo, G. Sonder, J. R. Schwebke, E. Hoornenborg, R. W. Peeling, S. S. Philip, N. Low, and C. K. Fairley. 2017. Sexually transmitted infections: challenges ahead. Lancet Infect Dis 17:e235–e279.

4. World Health Organization (WHO). 2016. WHO guidelines for the treatment of Neisseria gonorrhoeae. WHO: Geneva, Switzerland, 2016; Available from: https://www.who.int/reproductivehealth/publications/rtis/gonorrhoea-treatment-guidelines/en/.

5. Fifer, H., J. Saunders, S. Soni, S. T. Sadiq, and M. FitzGerald. 2020. 2018 UK national guideline for the management of infection with Neisseria gonorrhoeae. Int J STD AIDS 31:4–15.

6. St, C. S., L. Barbee, K. A. Workowski, L. H. Bachmann, C. Pham, K. Schlanger, E. Torrone, H. Weinstock, E. N. Kersh, and P. Thorpe. 2020. Update to CDC’s Treatment Guidelines for Gonococcal Infection, 2020. MMWR Morb Mortal Wkly Rep 69:1911–1916.

7. Ohnishi, M., D. Golparian, K. Shimuta, T. Saika, S. Hoshina, K. Iwasaku, S. Nakayama, J. Kitawaki, and M. Unemo. 2011. Is Neisseria gonorrhoeae initiating a future era of untreatable gonorrheaã: detailed characterization of the first strain with high-level resistance to ceftriaxone. Antimicrob Agents Chemother 55:3538–45.

8. Camara, J., J. Serra, J. Ayats, T. Bastida, D. Carnicer-Pont, A. Andreu, and C. Ardanuy. 2012. Molecular characterization of two high-level ceftriaxone-resistant Neisseria gonorrhoeae isolates detected in Catalonia, Spain. J Antimicrob Chemother 67:1858–60.

9. Unemo, M., D. Golparian, R. Nicholas, M. Ohnishi, A. Gallay, and P. Sednaoui. 2012. High-level cefixime-and ceftriaxone-resistant Neisseria gonorrhoeae in France: novel penA mosaic allele in a successful international clone causes treatment failure. Antimicrob Agents Chemother 56:1273–80.

10. Fifer, H., U. Natarajan, L. Jones, S. Alexander, G. Hughes, D. Golparian, and M. Unemo. 2016. Failure of Dual Antimicrobial Therapy in Treatment of Gonorrhea. N Engl J Med 374:2504–6.

11. Eyre, D. W., N. D. Sanderson, E. Lord, N. Regisford-Reimmer, K. Chau, L. Barker, M. Morgan, R. Newnham, D. Golparian, M. Unemo, D. W. Crook, T. E. Peto, G. Hughes, M. J. Cole, H. Fifer, A. Edwards, and M. I. Andersson. 2018. Gonorrhoea treatment failure caused by a Neisseria gonorrhoeae strain with combined ceftriaxone and high-level azithromycin resistance, England, February 2018. Euro Surveill 23.

12. Ross, J. D., and D. A. Lewis. 2012. Cephalosporin resistant Neisseria gonorrhoeae: time to consider gentamicinã Sex Transm Infect 88:6–8.

13. Chisholm, S. A., N. Quaye, M. J. Cole, H. Fredlund, S. Hoffmann, J. S. Jensen, M. J. van de Laar, M. Unemo, and C. A. Ison. 2011. An evaluation of gentamicin susceptibility of Neisseria gonorrhoeae isolates in Europe. J Antimicrob Chemother 66:592–595.

14. Lagace-Wiens, P., H. J. Adam, N. M. Laing, M. R. Baxter, I. Martin, M. R. Mulvey, J. A. Karlowsky, D. J. Hoban, and G. G. Zhanel. 2017. Antimicrobial susceptibility of clinical isolates of Neisseria gonorrhoeae to alternative antimicrobials with therapeutic potential. J Antimicrob Chemother 72:2273–2277.

15. Bala, M., V. Singh, A. Bhargava, M. Kakran, N. C. Joshi, and R. Bhatnagar. 2016. Gentamicin Susceptibility among a Sample of Multidrug-Resistant Neisseria gonorrhoeae Isolates in India. Antimicrob Agents Chemother 60:7518–7521.

16. Mann, L. M., R. D. Kirkcaldy, J. R. Papp, and E. A. Torrone. 2018. Susceptibility of Neisseria gonorrhoeae to Gentamicin—Gonococcal Isolate Surveillance Project, 2015-2016. Sex Trans Dis 45:96–98.

17. Lule, G., F. M. Behets, I. F. Hoffman, G. Dallabetta, H. A. Hamilton, S. Moeng, G. Liomba, and M. S. Cohen. 1994. STD/HIV control in Malawi and the search for affordable and effective urethritis therapy: a first field evaluation. Genitourin Med 70:384–388.

18. Kirkcaldy, R. D., H. S. Weinstock, P. C. Moore, S. S. Philip, H. C. Wiesenfeld, J. R. Papp, P. R. Kerndt, S. Johnson, K. G. Ghanem, E. W. Hook, L. M. Newman, D. Dowell, C. Deal, J. Glock, L. Venkatasubramanian, L. McNeil, C. Perlowski, J. Y. Lee, S. Lensing, N. Trainor, S. Fuller, A. Herrera, J. S. Carlson, H. Harbison, C. Lenderman, P. Dixon, A. Whittington, I. Macio, C. Priest, A. Jett, T. Campbell, A. Uniyal, L. Royal, M. Mejia, J. Vonghack, S. Tobias, J. Zenilman, J. Long, A. Harvey, K. Pettus, and S. Sharpe. 2014. The Efficacy and Safety of Gentamicin Plus Azithromycin and Gemifloxacin Plus Azithromycin as Treatment of Uncomplicated Gonorrhea. Clin Infect Dis 59:1083–1091.

19. Ross, J. D., J. Harding, L. Duley, A. A. Montgomery, T. Hepburn, W. Tan, C. Brittain, G. Meakin, K. Sprange, S. Thandi, L. Jackson, T. Roberts, J. Wilson, J. White, C. Dewsnap, M. Cole, and T. Lawrence. 2019. Gentamicin as an alternative to ceftriaxone in the treatment of gonorrhoea: the G-TOG non-inferiority RCT. Health Technol Assess 23:1–104.

20. Bollenbach, T. 2015. Antimicrobial interactions: mechanisms and implications for drug discovery and resistance evolution. Curr Opin Microbiol 27:1–9.

21. Yang, F., J. Yan, J. Zhang, and S. van der Veen. 2020. Evaluation of alternative antibiotics for susceptibility of gonococcal isolates from China. Int J Antimicrob Agents 55:105846.

22. Liu, J. W., W. Q. Xu, X. Y. Zhu, X. Q. Dai, S. C. Chen, Y. Han, J. Liu, X. S. Chen, and Y. P. Yin. 2019. Gentamicin susceptibility of Neisseria gonorrhoeae isolates from 7 provinces in China. Infect Drug Resist 12:2471–2476.

23. Daly, C. C., I. Hoffman, M. Hobbs, M. Maida, D. Zimba, R. Davis, G. Mughogho, and M. S. Cohen. 1997. Development of an antimicrobial susceptibility surveillance system for Neisseria gonorrhoeae in Malawi: comparison of methods. J Clin Microbiol 35:2985–8.

24. Lee, H., K. Lee, and Y. Chong. 2016. New treatment options for infections caused by increasingly antimicrobial-resistant Neisseria gonorrhoeae. Expert Rev Anti Infect ther 14:243.

25. White, R. L., D. S. Burgess, M. Manduru, and J. A. Bosso. 1996. Comparison of three different in vitro methods of detecting synergy: time-kill, checkerboard, and E test. Antimicrob Agents Chemother 40:1914–8.

26. Barbee, L. A., O. O. Soge, K. K. Holmes, and M. R. Golden. 2014. In vitro synergy testing of novel antimicrobial combination therapies against Neisseria gonorrhoeae. J Antimicrob Chemother 69:1572–1578.

27. Wind, C. M., H. J. de Vries, and A. P. van Dam. 2015. Determination of in vitro synergy for dual antimicrobial therapy against resistant Neisseria gonorrhoeae using Etest and agar dilution. Int J Antimicrob Agents 45:305–8.

28. Singh, V., M. Bala, A. Bhargava, M. Kakran, and R. Bhatnagar. 2018. In vitro efficacy of 21 dual antimicrobial combinations comprising novel and currently recommended combinations for treatment of drug resistant gonorrhoea in future era. PLoS One 13:e0193678.

29. Furuya, R., Y. Koga, S. Irie, M. Tanaka, F. Ikeda, A. Kanayama, and I. Kobayashi. 2013. In vitro activities of antimicrobial combinations against clinical isolates of Neisseria gonorrhoeae. J Infect Chemother 19:1218–1220.

30. Bharat, A., I. Martin, G. G. Zhanel, and M. R. Mulvey. 2016. In vitro potency and combination testing of antimicrobial agents against Neisseria gonorrhoeae. J Infect Chemother 22:194–7.

31. Unemo, M., D. Golparian, A. Limnios, D. Whiley, M. Ohnishi, M. M. Lahra, and J. W. Tapsall. 2012. In vitro activity of ertapenem versus ceftriaxone against Neisseria gonorrhoeae isolates with highly diverse ceftriaxone MIC values and effects of ceftriaxone resistance determinants: ertapenem for treatment of gonorrheaã Antimicrob Agents Chemother 56:3603–9.

32. Sood, S., S. K. Agarwal, R. Singh, S. Gupta, and V. K. Sharma. 2019. In vitro assessment of gentamicin and azithromycin-based combination therapy against Neisseria gonorrhoeae isolates in India. J Med Microbiol 68:555–559.

33. Pereira, R., M. J. Cole, and C. A. Ison. 2013. Combination therapy for gonorrhoea: in vitro synergy testing. J Antimicrob Chemother 68:640–643.

34. Siber, G. R., P. Echeverria, A. L. Smith, J. W. Paisley, and D. H. Smith. 1975. Pharmacokinetics of gentamicin in children and adults. J Infect Dis 132:637–51.

35. Richards, D. M., R. C. Heel, R. N. Brogden, T. M. Speight, and G. S. Avery. 1984. Ceftriaxone. A review of its antibacterial activity, pharmacological properties and therapeutic use. Drugs 27:469–527.

36. Nix, D. E., A. K. Majumdar, and M. J. DiNubile. 2004. Pharmacokinetics and pharmacodynamics of ertapenem: an overview for clinicians. J Antimicrob Chemother 53 Suppl 2:ii23–8.

37. Crokaert, F., A. Hubloux, and P. Cauchie. 1998. A Phase I Determination of Azithromycin in Plasma during a 6-Week Period in Normal Volunteers after a Standard Dose of 500mg Once Daily for 3 Days. Clin Drug Investig 16:161–6.

38. Clinical and Laboratory Standards Institute. 2018. Performance Standards for Antimicrobial Susceptibility Testing: Twenty-Eighth Informational Supplement M100. CLSI, Wayne, PA, USA.

39. The European Committee on Antimicrobial Susceptibility Testing. 2015. Breakpoint tables for interpretation of MICs and zone diameters. Version 5.0, 2015. https://www.eucast.org/fileadmin/src/media/PDFs/EUCAST_files/Breakpoint_tables/v_5.0_Breakpoint_Table_01.pdf.

40. WHO. 2012. Global Action Plan to Control the Spread and Impact of Antimicrobial Resistance in Neisseria gonorrhoeae. Available from:http://whqlibdoc.who.int/publications/2012/9789241503501_eng.pdf?ua=1

41. Tapsall, J. W., F. Ndowa, D. A. Lewis, and M. Unemo. 2009. Meeting the public health challenge of multidrug-and extensively drug-resistant Neisseria gonorrhoeae. Expert Rev Anti Infect Ther 7:821–34.

